# *CUT&ID* for simultaneous profiling of protein-DNA and protein-protein interactions

**DOI:** 10.64898/2025.12.11.693662

**Authors:** Anna Nordin, Claudio Cantù

## Abstract

We developed CUT&ID to simultaneously profile the native genomics and proteomics associations of any endogenous target, in a single streamlined workflow with no need of transgenesis. CUT&ID is enabled by a fusion protein that drives sequential target recognition, proximal protein biotinylation and DNA cleavage in living cells. CUT&ID is broadly applicable and yields high signal-to-noise genomics and proteomics identification across all types of gene regulators tested.

## Introduction

Gene expression drives cell fate specification and cell identity, and its dysregulation is a hallmark of essentially every disease. This process involves a complex system of transcription factors (TFs) and co-factors that, in addition to associating to genomic regulatory regions, work together to activate or inhibit transcription ^1^. Both these aspects are crucial to understanding gene regulation. Recent advancements have led to methods for analyzing either the genomic binding or the protein-protein interactions of a given gene regulator ^2–9^. While these have allowed for lower sample inputs and reduced background noise, the two scopes (identifying the interactions with DNA or with other proteins) have largely remained separated and require different methods. Here we develop CUT&ID, a method that allows the simultaneous profiling of both the DNA binding and protein partners of all classes of gene regulators (histone marks, transcription factors, non-DNA binding proteins), without crosslinking, tagging, and with modest inputs (1 million cells). The CUT&ID workflow is performed in three days, allows multi-channel pipette scale-up for increased throughput, and is successful across the board of all gene regulators tested.

## Results

We designed a fusion protein containing protein A/G (pAG) for antibody recognition, Micrococcal Nuclease (MNase) for DNA under target cleavage, and TurboID for proximal protein biotinylation. We first used AlphaFold ^10^ to model different possible configurations and chose Nter-TurboID-pAG-MNase-Cter (hereafter referred to as *Lazarus*), carrying two intervening flexible polypeptide linkers to ensure proper folding and functionality of all domains (Fig. 1A). *Lazarus* can be expressed in bacteria and purified via standard protocols. It displays robust enzymatic activities *in vitro*: MNase cleaves DNA upon addition of CaCl_2_ (Fig. 1B, left), and TurboID biotinylates itself and other surrounding proteins upon addition of biotin and ATP (Fig. 1B, right).

**Figure 1.**
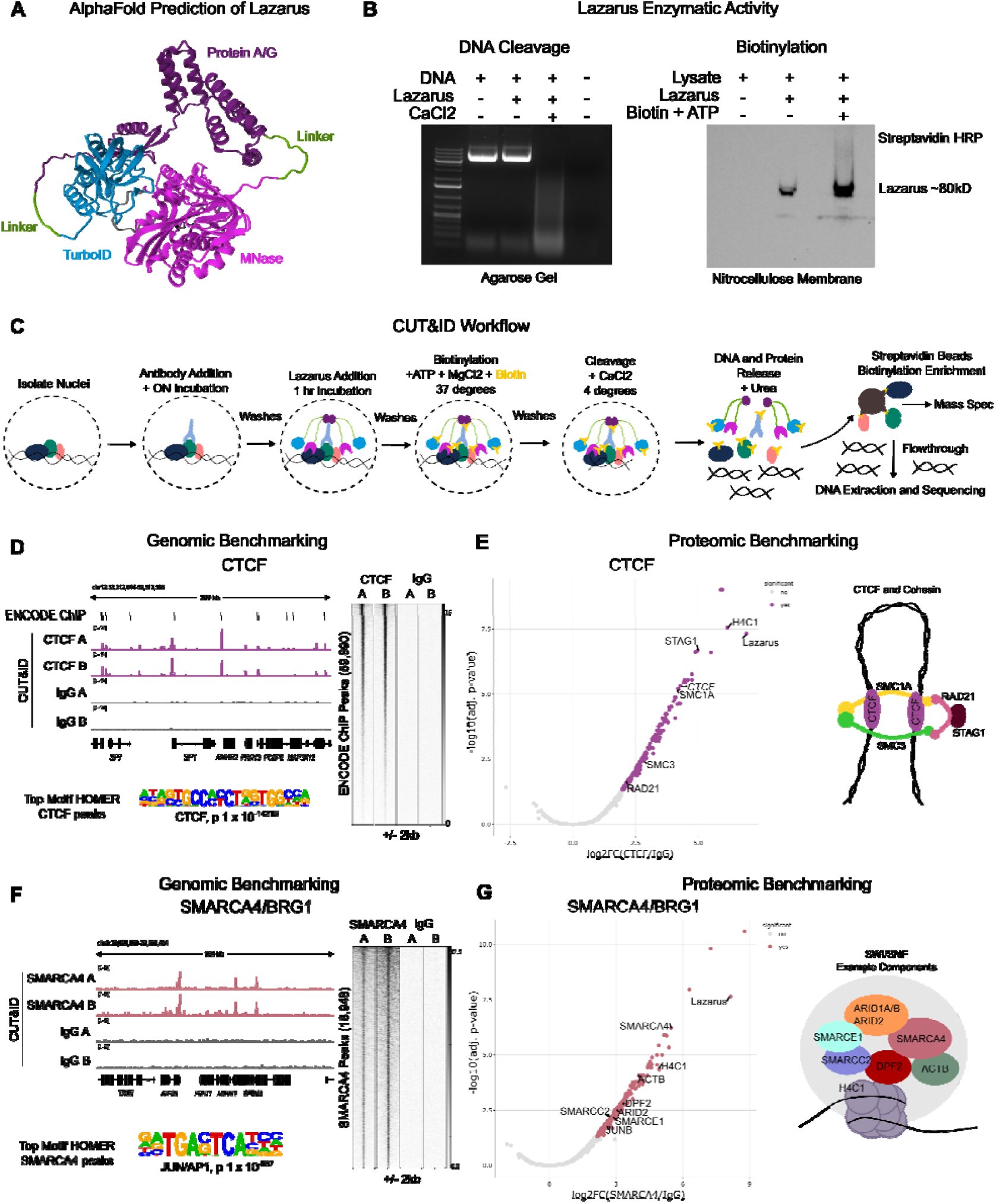
CUT&ID workflow and benchmarking. **A**. AlphaFold predicted structure of Nter-TurboID-pAG-MNase-Cter recombinant protein (referred to as Lazarus), Lazarus, containing from N to C termini: TurboID (blue), linker (green), protein A/G (purple), linker (green) and MNase (pink). **B**. Enzymatic in vitro assays showing that Lazarus cleaves DNA upon CaCl_2_ addition (left, DNA agarose gel) and biotinylates proteins in a lysate after addition of biotin and ATP (right, streptavidin-HRP western blot). **C**. Schematic depiction of the CUT&ID workflow. **D**. Genomic benchmarking of CTCF, showing IGV browser tracks of CUT&ID for CTCF, IgG and ChIP-seq peaks (top), top enriched HOMER motif (bottom), and high signal within ENCODE ChIP-seq peaks (right). **E**. Proteomic benchmarking of CTCF represented as volcano plot of CTCF/IgG enrichment (left) and the recovery of ‘textbook’ interactors. **F**. Genomic benchmarking of SMARCA4, showing genome browser view (top), top enriched motif (bottom) and signal enrichment within called peaks (right). **G**. Proteomic benchmarking of SMARCA4, showing volcano plot of SMARCA4/IgG enrichment (left) and the recovery of ‘textbook’ interactors. Peaks were called on merged replicates with MACS2 q < 0.05, and proteomic significance calculated using amica on default settings, with Log_2_FC > 1, adj. p < 0.05. Genomic datasets were normalized by reads per genome coverage and group scaled.

We optimized the CUT&ID workflow via gradual tinkering starting from combined CUT&RUN LoV-U ^5^ and TurboID ^7^ protocols (Fig. 1C; detailed step-by-step protocol in Supp. File 1). We first benchmarked CUT&ID on the two broadly relevant genome regulators: CTCF and SMARCA4/BRG1. The genomics data of individual replicates were merged after quality control, and peaks called with MACS2 against the IgG negative controls. The proteomic data was processed with Spectronaut for identification, quantification, and normalization, and then with amica ^11^ on default settings for imputation and differential enrichment analysis. Peaks are listed in Supp. Table 1, and enriched proteins in Supp. Table 2.

The CTCF genomics datasets exhibit the hallmarks of a successful CUT&RUN: high signal to noise ratio (FrIP scores 0.12 and 0.14, 34,941 called peaks at MACS2 q < 0.05), enrichment of the primary motif (CTCF, p 1 × 10^-14216^), and a high degree of overlap with ENCODE ChIP-seq data ^12^ (71% of CUT&ID peaks shared) (Fig. 1D). Quality control (QC) metrics of all genomic datasets are in Fig. S1A-D. In the proteomics part we identified 202 enriched proteins over the IgG control (Log_2_FC > 1, adj. p < 0.05), including CTCF itself (Log_2_FC 3.9, adj. p 3.0 x 10^-5^) and known interactors like STAG1, SMC1A, SMC3, and RAD21 (Fig. 1E, left) – all cohesin subunits (Fig. 1E, right) ^13^. Fold changes were calculated based on imputed values because in the IgG samples many of these proteins were not detected at all—a fact that supports the sensitivity of CUT&ID. Identified proteins, imputed intensities, and QC metrics of all proteomics are detailed in Fig S2 A-D. The recent method PLAM-seq, applied to CTCF, filtered by us with the same thresholds, yielded 28 enriched proteins overlapping with CUT&ID, including many ‘text-book’ interactors (Fig. 1E, right) ^9^.

The SMARCA4 experiment was similarly successful, with high signal-to-noise ratios for both genomics and proteomics identifications (Fig. 1F-G). We identified 18,948 genomic binding sites (MACS2 q < 0.05), with JUN/AP1 as the highest enriched consensus motif (Fig. 1F), and 219 enriched proteins, including SMARCA4 (Log_2_FC 5.2, adj. p 1.3 x 10^-6^), JUN/AP1, and other members of the SWI/SNF complex (ARID2, SMARCC2, SMARCE1, and DPF2) (Fig. 1F) ^14^. Also ProtA-Turbo ^8^, but not IgG control samples, identified SWI/SNF complex members. These results show that CUT&ID is on par with individual state-of-the-art methods for genomics (CUT&RUN, ChIP-seq) and proteomics (PLAM-seq, ProtA-Turbo), but yields dual omics readouts in one shot from lower input requirements, with no sample splitting, and no need of endogenous tagging or overexpression.

We tested CUT&ID for 6 additional targets to cover the spectrum of possible chromatin-relevant targets: chromatin marks (H3K27ac), transcription factors (TCF7L2 and JUN), and several challenging non-DNA binding co-factors (β-catenin, RAD21, and HDAC1). CUT&ID worked for all targets, yielding robust genomic and proteomic datasets that align well with existing data and known attributes for the targets (Fig. 2A-F). H3K27ac gave 17,445 genomic peaks and 301 enriched proximal proteins, where prominent were histones H3-7 (Log_2_FC 5.8, adj. p 2.5 × 10^-3^) and many chromatin regulators (Fig. 2A). TCF7L2 yielded in 24,273 genomic peaks and 36 high-confidence protein interactors, among which at the top were TCF7L2 (Log_2_FC 4.5, adj. p 4.5 × 10^-3^), β-catenin, and JUNB. Genomics and proteomics identifications were in line with current knowledge ^15^ and with each another—TCF7L2 and JUNB protein identification matched the enriched consensus motifs for TCF/LEF (p 1 × 10^-502^) and JUNB (p 1 × 10^-1836^) (Fig. 2B). β-catenin was found to occupy 6282 genomic regions, of which 5970 (95%) overlapped with those bound by TCF7L2, as expected based on Wnt signaling mechanisms ^15^. The top proteomic hit was β-catenin itself (Log_2_FC 6.1, adj. p 6.9 × 10^-5^) together with the members of the adherens junctions α-catenin and CDH1 (Fig. 2C). For JUN we obtained 4239 genomic peaks (top motif JUN/AP1 p 1 × 10^-204^), and 890 enriched proteins, with top hits of JUND (Log_2_FC 4.5, adj. p 2.6 × 10^-3^) and JUNB (Log_2_FC 3.8, adj. p 2.6 × 10^-3^), as well as reciprocal enrichment of TCF7L2 (Fig. 2D), cross-validating the TCF7L2 experiment. RAD21 resulted in 6133 genomic peaks (top motif CTCF, p 1 × 10^-2274^) and closely matched the CTCF datasets (Fig. S1D). Note that we previously had to perform crosslinking to obtain RAD21 CUT&RUN data ^16^ : CUT&ID did not require cross-linking even to detect this elusive target. We identified 51 high-confidence protein partners, including RAD21 (Log_2_FC 3.8, adj. 1.8 × 10^-3^), and other known interactors like STAG1, STAG2, SMC1A, and SMC3 (Fig. 2E) ^13^. Lastly, we obtained 11,466 HDAC1 peaks and 614 vicinal proteins, including HDAC1 (Log_2_FC 4.3, adj. p 5.8 × 10^-4^), and members of histone de-acetylation complexes such as SIN3A and DNTTIP1 ^17^. Taken together, these results demonstrate that CUT&ID is widely applicable to diverse targets, obtaining highly specific genomic and proteomic profiles.

**Figure 2.**
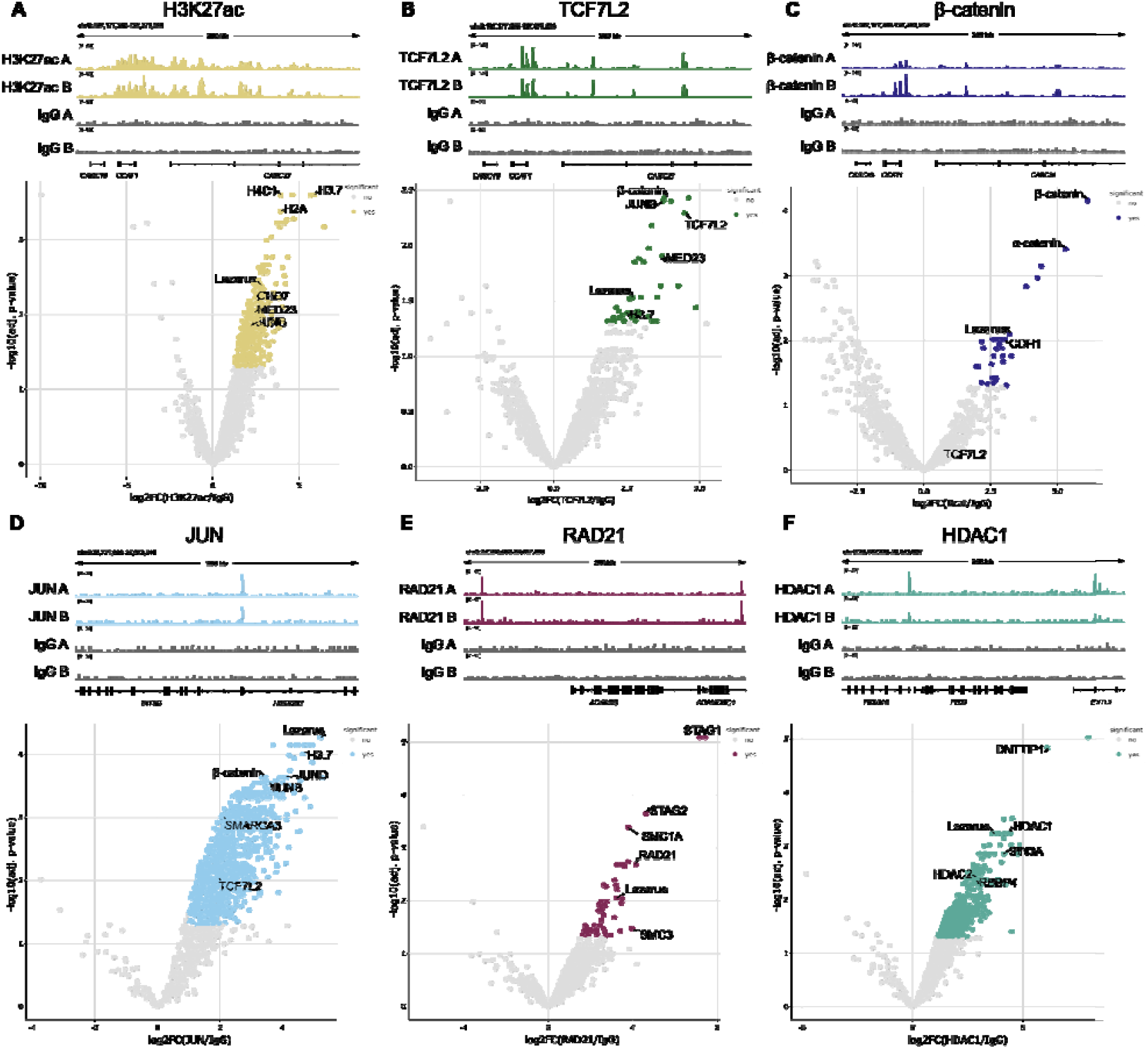
Applying CUT&ID to gene regulators. IGV Browser view (top) and volcano enrichment of proteins (bottom) using CUT&ID H3K27ac, (**B**) TCF7L2 (**C**) β-catenin, (**D**) JUN, (**E**) RAD21, (**F**) HDAC1. Examples of known interactors are labelled. Peaks were called on merged replicates with MACS2 q < 0.05, and proteomic significance calculated using amica on default settings, with Log_2_FC > 1, adj. p < 0.05. Genomic datasets were normalized by reads per genome coverage and group scaled.

## Discussion

CUT&ID emerges as a versatile and simple method to simultaneously map DNA binding and proximal protein interactions from the same sample and procedure. CUT&ID is robust, reproducible, and broadly applicable across histone modifications, TFs, and difficult to detect non-DNA binding cofactors. We observed strong recovery of expected genomic features, including high signal-to-noise ratios, appropriate motif enrichments, and overlap with reference datasets, confirming that CUT&ID performs indistinguishably from CUT&RUN. In parallel, protein proximity labeling reliably identifies known interactors, with the bait protein invariably enriched at the top of the list, a necessary internal control of selective identification. The dual readout of CUT&ID also provides a unique opportunity to evaluate antibody specificity: proteins identified by mass spectrometry provide an orthogonal quality check for the genomic dataset, revealing whether off-target antibody binding might contribute to unexpected peaks. This built-in validation increases confidence in both the antibody and the specificity of the resulting datasets, which is an important consideration for the gene regulation community.

CUT&ID has several advantages. It does not require cloning and overexpression nor endogenous knock-ins (as in BioID/TurboID ^7^ or PLAM-seq ^9^), making it readily employable for any endogenous targets across models, *ex vivo* tissues, and even patients’ samples. It does not require crosslinking of cells or nuclei (like in ChIP-seq ^2^, PLAM-seq ^9^ and RIME ^6^), reducing the introduction of undesired artefacts and minimizing the variability across experiments caused by differences in fixation. Finally, it uses low input material, comparably to CUT&RUN ^3^, and less than ChIP-seq ^2^ or ProtA-Turbo ^8^— which requires 15 million cells, double if a ChIP-seq is to be performed in parallel. We used 1 million cells per sample yielding robust genomic and proteomic datasets across all targets tested. However, optimal input may vary depending on abundance, antibody affinity, and mass spectrometer sensitivity. For high-abundance proteins, or when using efficient antibodies, CUT&ID might be amenable to scaling down, while increasing input could enable the detection of more rare or transient interactions. We recommend target-specific optimization based on these parameters. CUT&ID is in fact designed with scalability in mind. The workflow uses low volumes, multichannel pipetting, and is compatible with parallel processing of dozens of samples and is easily adaptable to 96-well plates or automation.

## Acknowledgments

We are grateful to all the members of the Cantù lab for suggestions and discussions; to Tamina Weiss, Yorick van de Grift, Jente Zijlstra and Jakob Hajman for detailed comments on the manuscript; to Mattias Jonasson who conceived pAG-TID, a project subsequently abandoned as scooped by Santos-Barriopedro et al.; to former member Pierfrancesco Pagella who coined the name “*Lazarus*” — this project arose from the dead with the intuition from A.N. and C.C. to add MNase. We acknowledge the Core Facility at the Faculty of Medicine and Health Sciences, Linköping University for providing assistance in Mass Spectrometry and DNA sequencing. The computations and data handling were enabled by resources provided by the National Supercomputer Centre (NSC), funded by Linköping University. We thank Peter Münger at the NSC for assistance concerning technical and implementational aspects in making the codes run on the Sigma resource.

## Author contributions

A.N. and C.C. conceived the project. A.N. performed experiments, formal analysis, and prepared figures. C.C. supervised the research and provided financial support for the study. A.N. wrote the manuscript, CC revised it.

## Data availability

Raw sequencing data and bigwigs for visualization have been deposited at ArrayExpress (E-MTAB-16382). Raw proteomics data has been deposited at ProteomeXchange (PXD071610). Called peaks and enriched proteins are provided in Supp. Tables 1 and 2.

## Methods

### AlphaFold modeling

Structural predictions were done with ColabFold (v1.5.3) ^18^, with default settings and visualized and annotated using the Mol* Viewer software ^19^.

#### *Lazarus* protein production

The codon-optimized sequence for TurboID-pAG-MNase (*Lazarus*) was synthesized by GenScript and cloned into the pET-28a(+)-TEV vector and then transformed into BL21 (DE3) (Cat. #70235, Sigma Aldrich) *e. coli* cells and plated on kanamycin agar plates ON. Colonies were picked and grown first for 16 hours in 10 ml of LB broth, then 1 ml was taken and added to 200 ml of LB and grown until the OD reached 0.4. IPTG was added to a final concentration of 100 µM and the protein produced for 20 hours at RT (20 °C) shaking. Bacteria were pelleted at 3000 x g at 4°C for 15 min and then washed twice in 20 ml ice-cold PBS. Bacteria were resuspended in 20 ml 2X lysis buffer (300 mM NaCl, 20 mM Tris-HCl pH 7.4, 40 mM imidazole, 0.1 mM DTT, 0.005% IGEPAL, 0.1 mM PMSF) and resuspended by vortexing. Lysis was completed by sonication for 5 min in a tip-sonicator, 5 sec ON/OFF, amplitude 10. Lysate was cleared by centrifugation at max speed for 15 min at 4 °C. 250 µl of HisPur Ni-NTA resin (Cat. #88221, Thermo Scientific) was washed in 1X lysis buffer twice and then added to the lysate and incubated for 30 min in ice. The resin was pelleted by centrifugation at 500 x g for 2 min and then washed twice in 20 ml 1X lysis buffer and then resuspended once in 500 µl 1X lysis buffer. 10 µl TEV protease (Cat. #P8118S, NEB) was added and incubation proceeded ON at 4°C on a rotator. After incubation, the supernatant was kept and mixed with 500 µl 100% glycerol for storage. The resin was resuspended in 500 µl 1X lysis buffer with 250 mM imidazole for elution and mixed with 500 µl 100% glycerol for storage. Cleaved and eluted fractions were quantified and visualized with SDS-PAGE, and the cleaved fraction was used for experiments.

### Enzymatic testing

DNA cleavage was assessed by combining plasmid DNA with either EDTA or CaCl_2_ and *Lazarus* protein in PBS, followed by incubation for 30 min at 37 °C for cleavage, and 5 min at 70 °C for inactivation. Results were visualized on a 1% agarose gel with SYBR safe stain and visualized on the ChemiDoc (BioRad). Protein biotinylation was tested by lysing HCT116 cells in RIPA buffer with sonication, and then adding *Lazarus* alone or *Lazarus* with biotin and ATP and incubating for 1 hour at 37 °C. Samples were combined with 2X Laemmli, boiled at 95 °C for 10 minutes and run on a 10% polyacrylamide gel during SDS-PAGE at 120V for 90 min. Proteins were transferred to a nitrocellulose membrane with wet transfer ON at 4 °C at 30 V, and then the membrane was blocked for 1 hour RT with 5% BSA in TBS 0.1% Tween (TBST). The membrane was washed thrice with TBST and then incubated with Streptavidin-HRP (Cat. #21130, Thermo Scientific) diluted 1:5000 for 1 hour at RT and then washed six times. Pierce ECL western blotting substrate (Cat. #32106, Thermo Scientific) was used for detection and visualized on the ChemiDoc (Bio-Rad).

### Cell Culture

HCT116 human male colorectal cancer cells were cultured in a 37 °C incubator in 5% CO2 with humidity. Culture medium consisted of high glucose Dulbecco’s Modified Eagle Medium (Cat. #41965039, Gibco) with 10 % fetal bovine serum (Cat. #26140079, Gibco) and 1X PenStrep (Cat. #15140148, Gibco). 1,000,000 cells/sample were harvested by incubation with Trypsin-EDTA (Cat. # 25200056, Gibco) for 5 minutes. The trypsin was quenched with culture media, and the cells were washed twice with DPBS (Cat. #14190094, Thermo Fisher Scientific).

### CUT&ID

Nuclear extraction was performed by three washes Nuclear Extraction (NE) buffer (20 mM HEPES-KOH pH-8.2, 10 mM KCl, 0.5 mM Spermidine, 0.05% IGEPAL [0.05%], 20% glycerol [20%], 0.1 mM PMSF). The nuclei were resuspended after the final wash in 1 ml wash buffer (20 mM HEPES pH 7.5, 150 mM NaCl, 0.5 mM Spermidine, 0.1 mM PMSF) per sample and bound to 40 µl Magnetic ConA beads (Cat. #93569, Cell Signaling Technology) equilibrated in provided binding buffer. Bead binding proceeded for 15 min at 4 °C, after which nuclei and beads were resuspended in 200 µl wash buffer per sample and split into PCR tubes. Beads were collected on the magnet and then resuspended in 200 µl EDTA wash buffer (wash buffer with 2 mM EDTA) and incubated at RT for 5 min. Samples were then directly resuspended in antibody buffer (wash buffer with 2 µg antibody, 2 mM EDTA, 0.025% digitonin, and 0.05% BSA) and incubated ON at 4 °C on a rotator. Antibodies used included anti-CTCF (abcam, ab188408), anti-SMARCA4 (antibodies-online, ABIN6991990), anti-H3K27ac (antibodies online, ABIN2668475), anti-TCF7L2 (Cell Signaling Technology, C48H11), anti-β-catenin (antibodies online, ABIN2855042), anti-JUN (antibodies-online, ABIN3020286), anti-RAD21 (antibodies-online, ABIN2856242), anti-HDAC1 (antibodies-online, ABIN2854776), and rabbit IgG isotype control (Invitrogen, #100500C).

After ON incubation samples were washed 5 times (wash buffer with 0.025% digitonin) and resuspended in 200 µl of *Lazarus* protein buffer (wash buffer with 480 ng *Lazarus*/sample) and incubated for 1 hour at 4 °C on a rotator. Next, samples were washed five times and then resuspended in 100 µl biotinylation buffer (wash buffer with 5 mM MgCl2, 5 mM biotin, and 2 mM ATP) and incubated at 37 °C for 30 min. Samples were washed twice to wash out excess biotin, chilled in ice for 5 min during the last wash, and then resuspended in 100 µl ice cold digestion buffer (wash buffer with 2 mM CaCl2). Digestion was allowed to proceed for 30 min at 4 °C. Digestion was stopped by the addition of STOP mix (1 µl 500 mM EDTA, 1 µl 500 mM EGTA, 4 µl 5M NaCl). Samples were incubated at 37 °C for 30 minutes, and then microfuged for 30 sec. The first release was transferred to new PCR tubes, and the beads resuspended in 50 µl 1X Urea STOP buffer (100 mM NaCl, 2 mM EDTA, 2 mM EGTA, 0.5% IGEPAL, 8.6 M Urea) and incubated for 30 min at 20 °C. After incubation, samples were microcentrifuged for 30 sec and then the supernatant combined with the first release and the beads discarded. 20 µl Pierce streptavidin beads (Cat. #88817, Thermo Scientific) were prepared per sample by washing 3 times in PBS and then resuspending in 50 µl PBS with 0.1 mM PMSF. Beads were added to each sample (final volume 200 ml) and incubated at RT for 1 hour on a rotator. The beads were collected on a magnet, and supernatant containing DNA and non-biotinylated proteins were processed for DNA extraction (see DNA sequencing) and the beads attached to biotinylated proteins processed for mass spectrometry (see mass spectrometry).

### DNA sequencing

Supernatant from Pierce beads was transferred to new tubes containing 2 µl 10% SDS and 2.5 µl proteinase K, and incubated at 50 °C for 1 hour. 1 volume of phenol:chloroform:isoamyl alcohol was added to each sample and vortexed for 30 sec. Samples were centrifuged at 15,000 x g for 15 min, and then the aqueous phase transferred to new tubes containing 1 µl glycogen, 20 µl 3M sodium acetate, and 500 µl 100% EtOH. Samples were incubated ON at -20 °C, centrifuged at 15,000 x g at 4 °C for 45 min, and the supernatant discarded. Pellets were washed once with 1 ml 70% EtOH, centrifuged again for 10 min, and then dried for 10 min at 50 °C. Pellets were resuspended in 15 µl 10 mM Tris-HCl pH 8.0. Library preparation was performed with the KAPA EvoPrep kit (Cat. #10096039001, Roche) and KAPA dual-index adapters according to manufacturer’s instructions with the following modifications: 0.5X volumes were used throughout the protocol, libraries were amplified for 13 PCR cycles, and samples were size-selected by cutting from 150 – 500 bp after running on 2% agarose E-gel EX gels (Cat. #G8142ST, Invitrogen) with the Thermo Fisher E-Gel System. Samples were sequenced with 50 bp pair-end reads on the Illumina NextSeq 2000 with the NextSeq 1000/2000 P2 XLEAP RGTSKit (100c) to approximately 10 million raw reads/sample.

### Genomic Data Analysis

Reads were trimmed with bbmap bbduk (version 38.18) ^20^ to remove adapters and poly [AT], [G] and [C] repeat sequences. Reads were aligned to the hg38 genome with bowtie2 (version 2.4.5) ^21^ with the options --local --very-sensitive-local --no-unal --no-mixed --no-discordant --phred33 --dovetail -I 0 -X 500. Samtools (version 1.11) ^22^ suite was used to remove duplicate and incorrectly paired reads. Bedtools (version 2.30.0) ^23^ was used to remove reads mapped to the CUT&RUN hg38 suspect list from bam files. Bigwigs were created for visualization using deepTools (version 3.5.1-0) ^24^ with the options -e and -RPCG. After quality control of individual datasets, biological duplicates were merged using samtools. Peaks were called on merged datasets using MACS2 ^25^ with the options - -keep-dup all, -f bampe and -q 0.05. Heatmaps, signal intensity plots, and QC plots were created with deepTools. Motif analysis was done using HOMER (version 4.11) ^26^ findMotifsGenome with the hg38 genome using -size 200. Peaks for ENCODE CTCF ChIP-seq in HCT116 were downloaded for comparison (ENCSR240PRQ) ^12^ . Genome browser images were created with IGV ^27^ with RPCG normalized bigwigs.

### Mass Spectrometry

Pierce streptavidin beads were washed 5 times in protein wash (PW) buffer (10 mM Tris-HCl pH 7.5, 150 mM NaCl, 0.1% sodium deoxycholate, 0.1% SDS, 1% Triton X, 1 mM EDTA, 0.5 mM EGTA), once in 1M KCl, once in 0.1M Na2CO3, twice in PW buffer, transferred to a new tube, once in 2M Urea in 50 mM Tris HCl pH 7.5, once in 50 mM Tris-HCl pH 7.5, transferred to a new tube, and then thrice in 50 mM ammonium bicarbonate (ABC). All wash volumes were 200 ml. Beads were resuspended a final time in 100 µl ABC and transferred to new low bind tubes, and then 0.4 µg of Pierce Trypsin protease (Cat. #90057, Thermo Scientific) was added per sample and incubated for 1 hour at 37 °C. The supernatant was transferred to a new tube, and the beads washed twice with 50 µl ABC and the supernatants pooled (final volume 200 ml). Beads were discarded. Samples were centrifuged at 15,000 x g for 10 min, and then 180 µl transferred to new tubes containing 10 µl of 100 mM DTT and incubated for 30 min at 37 °C. 26 µl of 100 mM iodoacetamide was added and samples incubated at RT for 45 min in the dark. An additional 0.5 µg of Trypsin was added and the samples incubated ON at 37 °C on a rotator. After ON incubation, samples were dried in a vacuum centrifuge, reconstituted in 40 µl resuspension solution (0.5% TFA 5% ACN) and desalted using Pierce C18 columns (Cat. #89870, Thermo Scientific) according to manufacturer’s instructions. After desalting, samples were dried in a vacuum centrifuge and reconstituted in 0.1% formic acid. LC-MS/MS analysis was performed using a High Resolution Trapped Ion Mobility timsTOF HT mass spectrometer coupled to a nanoElute 2 LC system via a CaptiveSpray ionization source (Bruker Daltonics). 200 ng of peptide mixture sample reconstituted in 0.1% FA was loaded on reverse-phase C18 HPLC column (PepSep XTREME with dimensions of 25 cm x 150 µm x 1.5 µm, Bruker Daltonics) and peptides separated in 45 min with a gradient of 0.1% formic acid in water (A) and 0.1% formic acid in acetonitrile (B) as follows: from 2% B to 17% B over 25 minutes; from 25% B to 37% B over 35 minutes; from 37 % B to 95% B over 45 minutes with a flow rate of 400 nl/min and a column temperature of 50°C. All data were acquired under the dia-PASEF mode with a MS1 scan range of 100-1700 m/z, and the collision energy was linearly interpolated between 1/K0 values, from 20 eV at 0.6 Vs/cm2 to 59 eV at 1.6 Vs/cm2, keeping constant above or below.

### Proteomics Data Analysis

dia-PASEF data files were analyzed in Spectronaut (version 20.2, Biognosys) using the human proteome FASTA with the addition of streptavidin, *Lazarus* enzyme and rabbit IgG heavy chain. The following parameters were applied for the search: trypsin as the proteolytic enzyme, allowing up to one missed cleavage; carbamidomethylation of cysteine as a fixed modification; oxidation of methionine as a variable modification, and minimum peptide length of 5. FDR was set to 0.05 and normalization applied with default settings. Protein quantities, run evidence count, and number of precursors were exported as a protein report, and then filtered for keratins, tubulins, and ribosomal proteins. For differential analysis, protein quantities were uploaded to amica ^11^ and processed with default settings. amica was used to visualize results and create figure panels. The mass spectrometry proteomics data have been deposited to the ProteomeXchange Consortium via the PRIDE partner repository with the dataset identifier PXD071610.

## Availability of Data and Materials

Raw sequencing data and bigwigs for visualization have been deposited at ArrayExpress (E-MTAB-16382). Raw proteomics data has been deposited at ProteomeXchange (PXD071610). Called peaks and enriched proteins are provided in Supp. Tables 1 and 2. Supp. File 1 contains a detailed, step-by step protocol for performing CUT&ID.

---

**Fig. S1.**
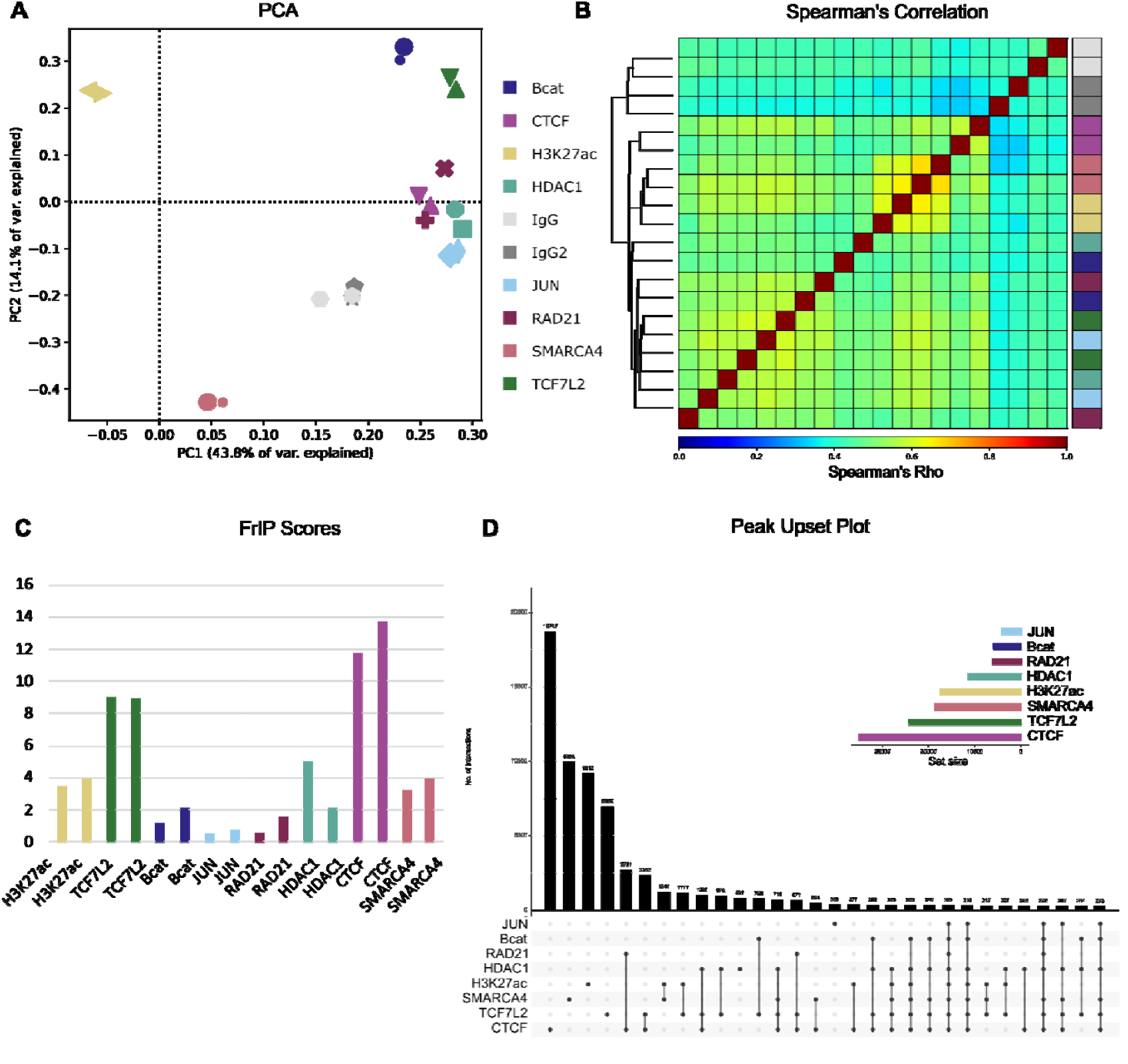
Quality control of genomic datasets. **A**. PCA of all samples done after genomic binning. IgG samples cluster together away from others **B**. Spearman’s correlation of all samples after genomic binning. **C**. FrIP scores of all replicates, based on peaks called on merged replicates (MACS2 q < 0.05). **D**. Upset plot of peak overlaps and peak set sizes. Expected overlaps between for example CTCF and RAD21, and TCF7L2 and β-catenin are seen.

**Fig. S2.**
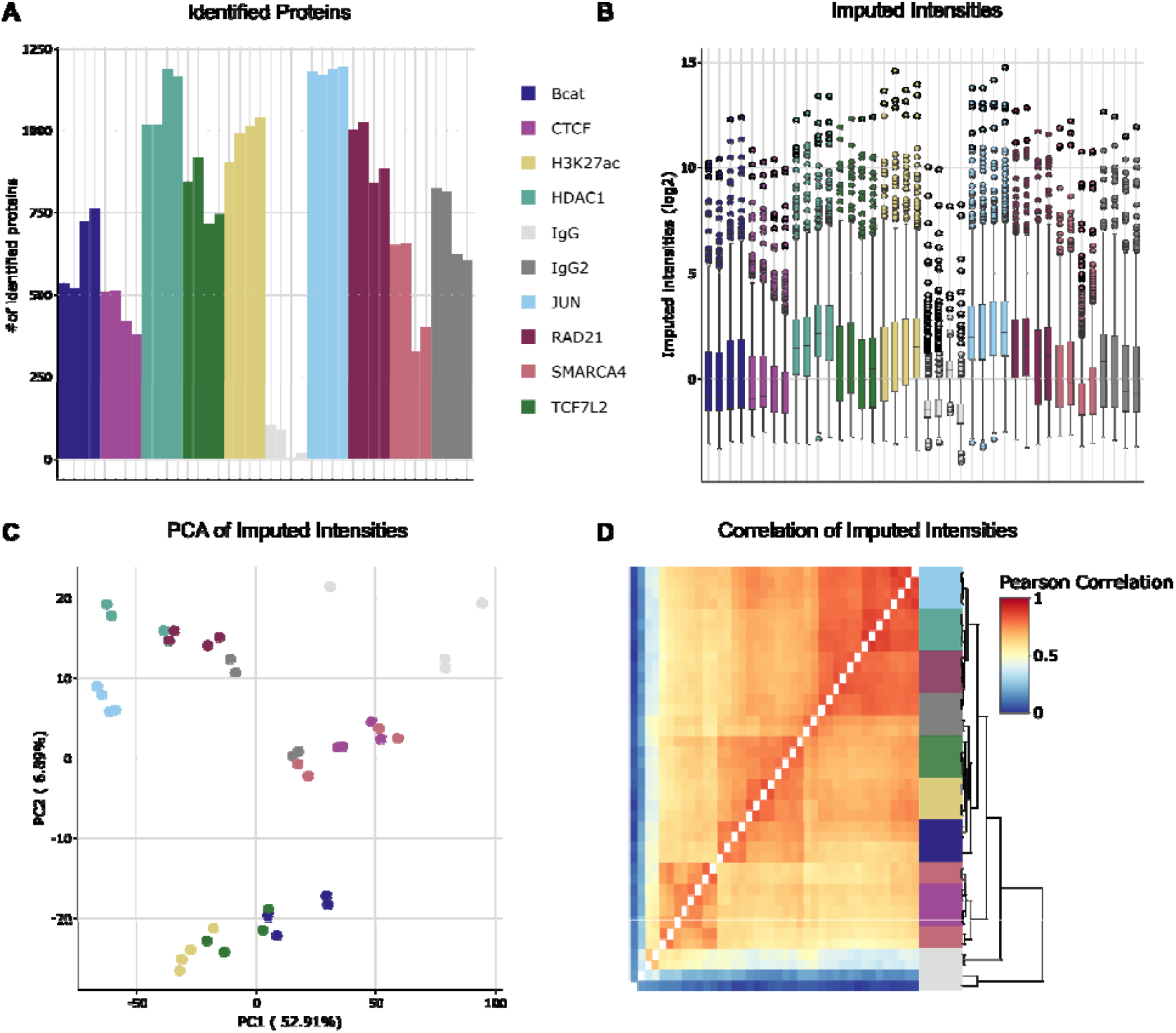
Quality control of proteomic datasets. **A**. Number of identified proteins in all replicates. CTCF and SMARCA4 were performed in parallel with IgG, and the rest with IgG2. Generally, the first experiment identified fewer proteins. **B**. Boxplot of imputed intensities for all samples. Imputation was performed with the amica software on default settings, after initial processing with Spectronaut. **C**. PCA of imputed intensities for all samples. **D**. Pearson’s correlation of imputed intensities of all samples.

## References

1. Lambert, S. A. et al. The Human Transcription Factors. Cell 172, 650–665 (2018).

2. Furey, T. S. ChIP–seq and beyond: new and improved methodologies to detect and characterize protein–DNA interactions. Nature Reviews Genetics 2012 13:12 13, 840–852 (2012).

3. Skene, P. J. & Henikoff, S. An efficient targeted nuclease strategy for high-resolution mapping of DNA binding sites. Elife 6, (2017).

4. Kaya-Okur, H. S. et al. CUT&Tag for efficient epigenomic profiling of small samples and single cells. Nat Commun 10, 1–10 (2019).

5. Zambanini, G., Nordin, A., Jonasson, M., Pagella, P. & Cantù, C. A new CUT&RUN low volume-urea (LoV-U) protocol optimized for transcriptional cofactors uncovers Wnt/β-catenin tissue-specific genomic targets. Development (Cambridge) 149, (2022).

6. Mohammed, H. et al. Rapid immunoprecipitation mass spectrometry of endogenous proteins (RIME) for analysis of chromatin complexes. Nat Protoc 11, 316–326 (2016).

7. Branon, T. C. et al. Efficient proximity labeling in living cells and organisms with TurboID. Nat Biotechnol 36, 880–898 (2018).

8. Santos-Barriopedro, I., van Mierlo, G. & Vermeulen, M. Off-the-shelf proximity biotinylation for interaction proteomics. Nat Commun 12, 1–12 (2021).

9. González-Vinceiro, L. et al. PLAMseq enables the proteo-genomic characterization of chromatin-associated proteins and protein interactions in a single workflow. Sci Adv 11, eady4151 (2025).

10. Jumper, J. et al. Highly accurate protein structure prediction with AlphaFold. Nature 2021 596:7873 596, 583–589 (2021).

11. Didusch, S., Madern, M., Hartl, M. & Baccarini, M. amica: an interactive and user-friendly web-platform for the analysis of proteomics data. BMC Genomics 23, (2022).

12. Dunham, I. et al. An integrated encyclopedia of DNA elements in the human genome. Nature 489, 57–74 (2012).

13. Merkenschlager, M. & Nora, E. P. CTCF and Cohesin in Genome Folding and Transcriptional Gene Regulation. Annu Rev Genomics Hum Genet 17, 17–43 (2016).

14. Nguyen, V. T., Tessema, M. & Weissman, B. E. The SWI/SNF Complex: A Frequently Mutated Chromatin Remodeling Complex in Cancer. Cancer Treat Res 190, 211–244 (2023).

15. Söderholm, S. & Cantù, C. The WNT/β-catenin dependent transcription: A tissue-specific business. WIREs mechanisms of disease 13, e1511 (2021).

16. Nordin, A. et al. Wnt signaling activation induces CTCF binding and loop formation at cis-regulatory elements of target genes. Genome Res 35, 1701–1716 (2025).

17. Adams, G. E., Chandru, A. & Cowley, S. M. Co-repressor, co-activator and general transcription factor: the many faces of the Sin3 histone deacetylase (HDAC) complex. Biochem J 475, 3921–3932 (2018).

18. Mirdita, M. et al. ColabFold: making protein folding accessible to all. Nature Methods 2022 19:6 19, 679–682 (2022).

19. Sehnal, D. et al. Mol* Viewer: modern web app for 3D visualization and analysis of large biomolecular structures. Nucleic Acids Res 49, W431–W437 (2021).

20. Bushnell, B., Rood, J. & Singer, E. BBMerge - Accurate paired shotgun read merging via overlap. PLoS One 12, (2017).

21. Langmead, B. & Salzberg, S. L. Fast gapped-read alignment with Bowtie 2. Nat Methods 9, 357 (2012).

22. Danecek, P. et al. Twelve years of SAMtools and BCFtools. Gigascience 10, 1–4 (2021).

23. Quinlan, A. R. & Hall, I. M. BEDTools: a flexible suite of utilities for comparing genomic features. Bioinformatics 26, 841–842 (2010).

24. Ramírez, F., Dündar, F., Diehl, S., Grüning, B. A. & Manke, T. deepTools: a flexible platform for exploring deep-sequencing data. Nucleic Acids Res 42, W187–W191 (2014).

25. Zhang, Y. et al. Model-based analysis of ChIP-Seq (MACS). Genome Biol 9, 1–9 (2008).

26. Heinz, S. et al. Simple Combinations of Lineage-Determining Transcription Factors Prime cis-Regulatory Elements Required for Macrophage and B Cell Identities. Mol Cell 38, 576–589 (2010).

27. Robinson, J. T. et al. Integrative genomics viewer. https://doi.org/10.1038/nbt0111-24 (2011) doi:10.1038/nbt0111-24.

