## Supplemental File 1 for "*CUT&ID* for simultaneous profiling of protein-DNA and protein-protein interactions"

**CUT&ID**

Cantù Lab

Updated December 2025

**Buffers CUT&ID**

*Make all buffers fresh daily and filter for best results!

**Nuclear Extraction Buffer (NE) (20 ml):**

| **Component** | **Final Conc.** | **Volume** |
| --- | --- | --- |
| 50% Glycerol | 20% | 8 ml |
| 1 M HEPES KOH pH 8.2 | 20 mM | 400 µl |
| 1 M KCl | 10 mM | 200 µl |
| 10% IGEPAL | 0.05% | 100 µl |
| 2 M Spermidine | 0.5 mM | 5 µl |
| 100 mM PMSF | 0.1 mM | 20 µl |
| H_2_O |  | to 20 ml |

**Binding Buffer (20 ml):**

| **Component** | **Final Conc.** | **Volume** |
| --- | --- | --- |
| 1 M HEPES pH 7.5 | 20 mM | 400 µl |
| 1 M MnCl_2_ | 1 mM | 20 µl |
| 2 M CaCl_2_ | 1 mM | 10 µl |
| 1 M KCl | 10 mM | 200 µl |
| H_2_O |  | to 20 ml |

**Wash Buffer (50 ml):**

| **Component** | **Final Conc.** | **Volume** |
| --- | --- | --- |
| 1 M HEPES pH 7.5 | 20 mM | 1 ml |
| 5 M NaCl | 150 mM | 1.5 ml |
| 2 M Spermidine | 0.5 mM | 12.5 µl |
| 100 mM PMSF | 0.1 mM | 50 µl |
| H_2_O |  | to 50 ml |
| 5% Digitonin (from day 2) | 0.025% | 250 µl |

**Antibody Buffer (2 ml):**

| **Component** | **Final Conc.** | **Volume** |
| --- | --- | --- |
| Wash Buffer |  | 1972 µl |
| 5% Digitonin | 0.025% | 10 µl |
| 0.5 M EDTA | 2 mM | 8 µl |
| 10% BSA | 0.05% | 10 µl |

**Biotin Buffer (2ml):**

| **Component** | **Final Conc.** | **Volume** |
| --- | --- | --- |
| Wash Buffer (with dig) | - | 1988 µl |
| 250 µM Biotin (make fresh) | 5 µM | 4 µl |
| 250 µM MgCl_2_ | 5 µM | 4 µl |
| 100 mM ATP | 2 mM | 4 µl |

**STOP Mix (per sample):**

| **Component** | **Final Conc.** | **Volume** |
| --- | --- | --- |
| 0.5 M EDTA | - | 1 µl |
| 0.5 M EGTA | - | 1 µl |
| 5 M NaCl | - | 4 µl |

**Urea STOP (2 ml):**

| **Component** | **Final Conc.** | **Volume** |
| --- | --- | --- |
| 8.8 M Urea | 8.5M | 1934 µl |
| 100% IGEPAL | 0.5% | 10 µl |
| 0.5 M EDTA | 2 mM | 8 µl |
| 0.5 M EGTA | 2 mM | 8 µl |
| 5 M NaCl | 100 mM | 40 µl |

**Protein Wash Buffer (PW) (20 ml):**

| **Component** | **Final Conc.** | **Volume** |
| --- | --- | --- |
| 1M Tris-HCl pH 7.5 | 10 mM | 200 µl |
| 5M NaCl | 150 mM | 600 µl |
| 10% sodium deoxycholate | 0.1% | 200 µl |
| 10% SDS | 0.1% | 200 µl |
| 10% Triton X | 1% | 2 ml |
| 0.5 M EDTA | 1 mM | 40 µl |
| 0.5 M EGTA | 0.5 mM | 20 µl |
| H_2_O |  | to 20 ml |

**Day 1 (approx. 1 - 2 hours, start in the afternoon):**

1. Harvest 1 million cells/sample. We recommend enzymatic digestion (Trypsin, TrypLE, etc.) making sure to quench well with media if needed. If multiple samples/replicates are coming from the same source, they can be processed in parallel until sample splitting (step 9), with up to 4 samples (4 mil. cells)/tube without adjusting volumes.

2. Centrifuge at an appropriate speed for cells (recommend 600 x g) and wash once in 2 ml PBS.

3. Centrifuge again and resuspend cells in 2 ml cold NE buffer. Pipette up and down well.

4. Centrifuge (800 x g) for 3 min, then discard supernatant and repeat NE wash until the cell pellet has become smaller and more translucent. We recommend 3 washes total.

5. Resuspend the nuclei in 1 ml wash buffer.

6. Prepare 40 µl ConA beads per sample according to instructions from the manufacturer, or by washing them twice in 400 µl (10X volume) of binding buffer. Resuspend in 40 µl binding buffer/sample.

7. Add appropriate amounts of the ConA bead slurry to nuclei suspension. Incubate for 15 min at 4 °C, or according to the manufacturer’s recommendations.

8. Place tubes on a magnet rack, remove supernatant, and wash twice with 200 µl wash buffer/sample by adding wash buffer, pipetting up and down, and then placing the tubes back on the magnet.

9. Split the samples into PCR strips and place them on the smaller magnet rack.

10. Dilute antibodies (2 µg/sample) in antibody buffer (200 µl/sample) and resuspend beads directly in the correct antibody buffer containing antibodies. Incubate ON ( ~16 hours) at 4 °C on a rotator.

**Day 2 (approx. 8 hours for 8 – 16 samples):**

1. Place PCR strips with samples on the magnet rack. Discard supernatant. Wash samples by resuspending the beads gently in 200 µl cold wash buffer (always with digitonin on day 2!). Pipette up and down 8 times.

2. Repeat step 1 for a total of 5 washes.

3. Discard supernatant and resuspend beads in 200 µl cold wash buffer containing 480 ng of Lazarus protein (pre-dilute Lazarus for the entire experiment in wash buffer) and incubate for 1 hour at 4 °C on a rotator.

4. Repeat step 1 for a total of 5 washes.

5. Discard supernatant and resuspend beads in 100 µl of biotin buffer. Incubate for 30 min at 37 °C (in a thermocycler, heated lid at 40 °C) for biotinylation of proximal proteins.

6. Repeat step 1 twice to wash out biotin and place PCR strips in ice for 5 min to chill.

7. Place back on the magnet, discard supernatant and resuspend beads in 100 µl wash buffer containing 2 mM CaCl_2_. Incubate for 30 min at 4 °C (thermocycler, heated lid off) for chromatin digestion.

8. Stop the reaction by adding 6 µl of the STOP mix and mix well by pipetting. Incubate for 30 min at 37 °C to start releasing chromatin (thermocycler, heated lid 40 °C).

9. Microcentrifuge samples for 30 sec and then place them on the magnet, transferring supernatant with the first release to new tubes. Place the first release on ice for now.

10. Resuspend beads in 100 µl of Urea STOP. Incubate at for 30 min at RT (20 °C in the thermocycler, heated lid off).

11. Microcentrifuge samples for 30 sec and then place them on the magnet. Transfer supernatant to combine it with the first release. Mix well.

*The soluble protein fraction and the cleaved DNA fragments are in the combined supernatant. Insoluble proteins and the genomic DNA remain on the ConA beads, these can be discarded. Proceed now only with the combined supernatants.

12. Prepare 20 µl Pierce streptavidin magnetic beads per sample by washing them 3 times in PBS (2 ml is enough for each wash for up to 16 samples). Resuspend them in 50 µl of PBS with 0.1 mM PMSF per sample.

13. Add Pierce beads to each sample and incubate 1 hr RT on a rotator.

14. Prepare 1.5 ml tubes for each sample with 2 µl 10% SDS and 2.5 µl proteinase K.

15. Place samples on the magnet. Transfer supernatant to the new tubes – this contains the DNA and the unbound (non-biotinylated) proteins. Mix well, and incubate for 1 hr at 50 °C.

*All bead washing steps below can be performed with a multichannel pipette. With different beads (not Pierce), beads may clump. Washing is optimized for the Pierce beads.

16. Resuspend streptavidin beads in 200 µl PW buffer and pipette 8 times to mix. Place samples back on the magnet and discard supernatant.

17. Repeat for a total of 5 washes. During washes 2 and 4, let the samples rotate 5 min at RT.

18. Place samples on the magnet and discard PW. Resuspend beads once in 1M KCl, and transfer to new tubes.

19. Place samples on the magnet and discard supernatant, resuspend once in 0.1 M sodium bicarbonate, discard supernatant, and then wash twice in PW buffer, the second wash rotating for 5 min at RT.

20. Discard PW buffer. Resuspend beads in 200 µl of 50 mM Tris HCl pH 7.5 with 2M Urea. Transfer to new tubes.

21. Resuspend beads in 200 µl 50 mM Tris HCl pH 7.5, discard supernatant, and then resuspend in 200 µl 50 mM ammonium bicarbonate (ABC). Transfer to new tubes.

*ABC should be prepared fresh, check that pH ~7.8.

22. Wash beads two more times with 200 µl ABC, then resuspend in 50 µl ABC.

23. Dilute 0.4 µg of Pierce Trypsin protease in 50 µl of ABC per sample, and aliquot into 1.5 ml protein low bind tubes.

24. Add the samples to the new low bind tubes (final volume 100 µl) and incubate for 1 hr at 37 °C on a rotator for on-bead digestion. This starts digesting the biotinylated proteins from the streptavidin beads, releasing them into the supernatant.

25. Return to the DNA samples. Add 1 volume (200 µl) phenol:chloroform:isoamyl alcohol to each sample and vortex for > 30 sec.

26. Centrifuge DNA samples at > 15,000 x g for 15 min. In the meantime, prepare new tubes with 1 µl 20 mg/ml glycogen, 20 µl 3M sodium acetate pH 5.2, and 500 µl 100% EtOH.

27. Carefully transfer the aqueous phase of each sample to the new tubes, invert several times to mix, and incubate ON at -20 °C.

28. After 1 hr incubation of the protein samples, place them on the magnet rack and move the supernatant to new tubes. Resuspend the beads twice in 50 µl ABC to collect residual protein, combining the supernatants each time (final volume 200 µl). Discard streptavidin beads.

29. Centrifuge the pooled supernatants at 15,000 x g for 10 min. In the meantime, prepare new low bind tubes with 10 µl of 100 mM DTT. Always make DTT fresh.

30. Transfer 180 µl of each sample to the new tubes. Incubate for 30 min at 37 °C.

31. Add 26 µl of 100 mM iodoacetamide to each sample and incubate 45 min RT in the dark. Always prepare iodoacetamide fresh and keep away from the light.

32. Add an additional 0.5 µg Trypsin to each sample, seal tubes with parafilm and incubate ON at 37 °C on a rotator.

**Day 3 (timing depends on speed vac):**

*Instructions here are for our setup, with C18 desalting spin columns. Presumably other methods (spin-tips, etc.) would work equally well.

1. Speed vacuum samples until dry (at 45 °C). You may see a small yellow, white or slightly brownish pellet.

2. Follow manufacturer's instructions for desalting. Resuspension is in 40 µl. Elution is done with 20 µl elution buffer 3 times, for a final volume of 60 µl.

3. Speed vacuum samples until dry (at 45 °C). Pellets should not be invisible, or nearly.

4. Samples are now ready to be reconstituted for mass spectrometry, or can be stored at -20 °C.

5. For DNA samples: centrifuge at > 15,000 x g at 4 °C for > 45 min.

6. Discard supernatant by decanting and then add 1 ml ice-cold 70% EtOH. Vortex until the pellet lifts from the bottom of the tube.

7. Centrifuge again for 10 min, and discard supernatant.

8. Dry DNA pellets at 50 °C for no more than 10 min. Resuspend in 20 µl Tris-HCl pH 7.5 or pH 8 for library preparation.

**Mass Spectrometry:**

1. To proceed with mass spec, resuspend samples in 20 µl 0.1% formic acid.

2. Sonicate for 5 min in a water bath sonicator, vortex for 30 sec, and then centrifuge at 13,000 x g for 10 min.

3. Quantify proteins on a Nanodrop and adjust concentrations to 0.1 mg/ml for LC/MS.

4. Load 10 µl into special mass spec tubes, careful not make bubbles.

**Library Preparation:**

Samples can be prepared for sequencing like any other CUT&RUN or ChIP samples, either with ligation or tagmentation based protocols. During size selection, take care to avoid primer-dimers.

For samples in the paper, we used the KAPA EvoPrep Kit with KAPA dual-index adapters, with 0.5X volume reactions, according to the instructions. We amplified for 13 cycles, and size selected for fragments between >120 and less than 500 bp (between 0 and 380 bp without adapters).

Samples should be sequenced pair-end, we have done 36 bp or 50 bp. We aim for 5 – 10 million reads per sample.
